# Placental transcriptome profiling in congenital Chagas disease: gene networks associated with transmission

**DOI:** 10.1101/2025.09.17.676751

**Authors:** Sofia Apodaca, Emiliano E. Campos, Kevin D. Calupiña, Carolina Davies, Silvia A. Longhi, Laura Kamenetzky, M. Paola Zago, Schijman Alejandro G.

## Abstract

Chagas disease, caused by *Trypanosoma cruzi*, affects over seven million people globally. Vertical transmission during pregnancy significantly contributes to the urban spread of the disease, even in non-endemic areas. The placental barrier plays a key role in preventing fetal infection, although the molecular mechanisms underlying congenital transmission remain unclear. To identify placental factors associated with transmission, we conducted a transcriptomic study comparing placental tissues from deliveries of congenitally infected (M+B+), exposed but uninfected (M+B−), and unexposed/uninfected (M−B−) newborns. Differential gene expression analysis of transmitting placentas revealed that *ENSG00000304767,* a novel lncRNA sense intronic to *CEMIP* was overexpressed as well as *CGB5*, while *CEMIP, CADM3*, *CADH11* and *PRXX1* were underexpressed. In non-transmitting placentas, the long non-coding RNA *MIR4300* was overexpressed while *CGB5* was underexpressed. These results suggest that cell adhesion and the extracellular matrix integrity are altered in transmitting placentas. Additionally, gene set enrichment analysis using the GO library revealed immune-related terms underrepresented in both infected mother groups, and confirmed that extracellular matrix processes, particularly collagen organization and metabolism, constitute important factors in transmission events. Analysis using the Cell Type library showed that extravillous trophoblasts were overrepresented in M+B+, but the opposite in M+B−. In contrast, syncytiotrophoblasts and villous cytotrophoblasts were overrepresented in non-transmitting vs. control cases. Immune-related placental cell types were consistently reduced in both M+ groups when compared to controls. The co-expression network analysis confirmed that the placental signaling and structural integrity were compromised in transmitting cases. *ENPP1* and *SLC16A10* emerged as hub genes with pivotal roles in the pathways altered during congenital infection. These findings highlight key placental transcriptional alterations linked to congenital *T. cruzi* transmission and provide insight into potential molecular mechanisms of fetal protection or susceptibility.

**Author summary:** Chagas disease, caused by the parasite *Trypanosoma cruzi*, can be passed from mother to baby during pregnancy, constituting the main transmission way in urban and non-endemic areas. The placenta normally acts as a barrier to protect the fetus, but how this barrier fails in congenitally transmitted cases is still not well understood. Thus, we analyzed the gene expression in placental tissues from three groups: infected mothers that transmitted the parasite, infected mothers that did not transmit it, and uninfected mothers. We found that certain genes involved in immune response, hormone regulation, and the structure of the placenta showed clear differences among these groups. In particular, *CGB5*, involved in the production of pregnancy hormones, showed opposite activity patterns depending on whether the mother transmitted the parasite or not. We also found that specific placental cell types that help anchor the placenta or form its protective outer layer changed their expression pattern in transmitting cases. These shifts suggest that both the physical structure and immune system of the placenta may be altered in a way that allows the parasite to reach the fetus. Our findings reveal important molecular clues that could help predict congenital transmission of Chagas disease.

## Introduction

Chagas disease (CD), caused by the protozoan parasite *Trypanosoma cruzi*, remains a major public health concern in Latin America and is increasingly recognized in non-endemic regions due to migration [1]. While vectorial and transfusional routes have declined in prevalence, mother-to-child transmission causing congenital Chagas disease (cCD) has emerged as the predominant mode of new infections in many areas, including urban and non-endemic settings. An estimated 5% of children born to infected mothers acquire the infection during pregnancy, yet the biological factors determining transmission risk remain poorly understood.

The placenta is the critical interface between maternal and fetal environments, acting as both a physical and immunological barrier against vertical transmission of pathogens [2, 3]. Despite its protective role, congenital transmission of *T. cruzi* still occurs, indicating that specific alterations in placental structure or function may facilitate the passage of the parasite to the fetus. Among the placental mechanisms to protect the fetus from the infection are fetal production of proinflammatory cytokines [4], fetal natural killer response [5] and epithelial trophoblast turnover [2,6–8]. However, little is known about the molecular and cellular changes in the placenta associated with *T. cruzi* transmission.

In general, it is proposed that in human diseases, on average, each gene interacts with four to eight other genes [9]. In this sense, gene expression networks are useful to identify hundreds of genes that are potentially interconnected, constituting a powerful tool not only to construct gene networks but also to detect gene modules and identify the central players (i.e., hub genes) within these modules.

Building on a previous study that examined gene expression in placentas from *T. cruzi*-infected mothers who did not transmit cCD versus non-infected mothers [10], the present study aimed to further characterize the placental gene expression profiles and gene co-expression networks. For the first time, we performed a transcriptomic analysis of placental tissues from three groups: infected newborns (M+B+), exposed but uninfected newborns (M+B−), and unexposed controls (M−B−). This integrative analysis provides novel insights into the host–parasite interaction at the maternal–fetal interface and identifies potential molecular markers involved in susceptibility to cCD.

## Results

### Enrollment of mother-infant pairs. Placental characterization

The characterization of the placental groups into transmitting, non-transmitting and negative controls were based on the Chagas disease diagnostic algorithm of mother/newborn pairs. In total, 192 pregnant women were enrolled; 162 of them were diagnosed as positive for Chagas Disease (M+) and 30 as negative (M-). When mothers tested positive, their babies were diagnosed, resulting in 12 positives (B+) and 92 negatives (B-). There were 58 newborns that did not complete the diagnostic algorithm (B0) and therefore were excluded from the study (Fig 1).

**Fig 1.**
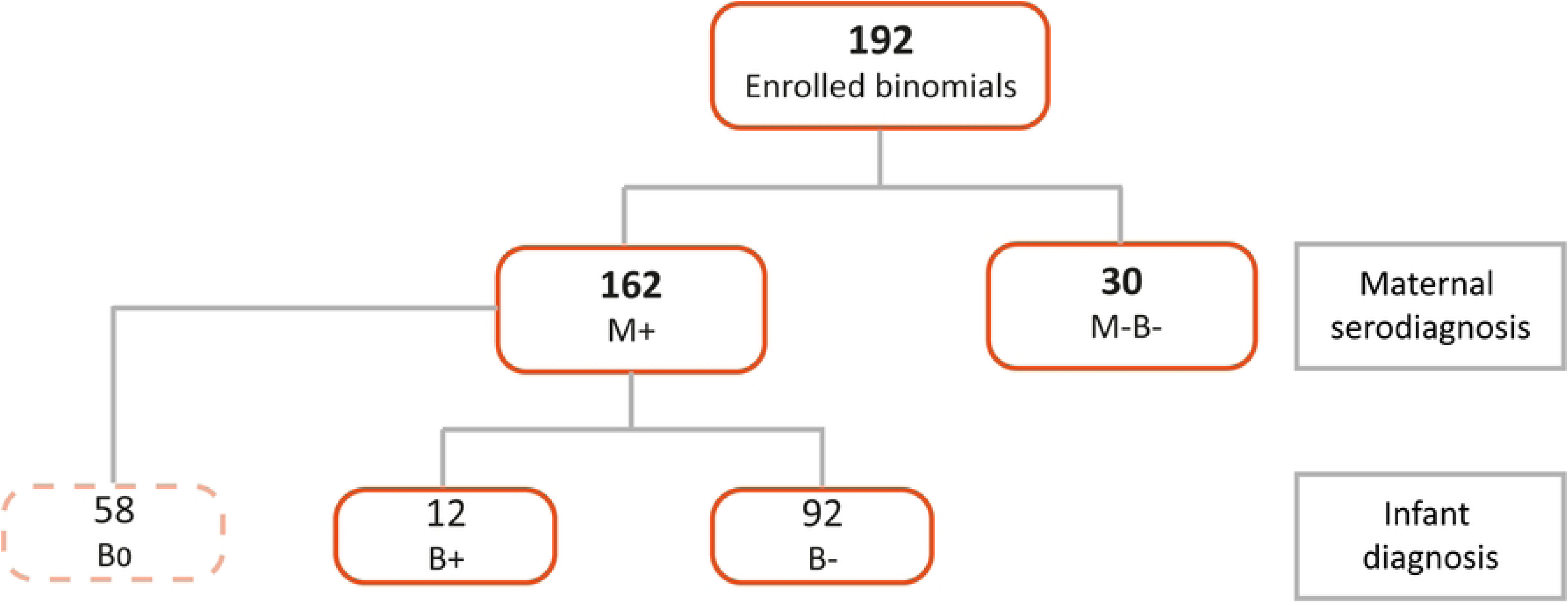
Flowchart of enrolled mother/newborn pairs to characterize placental groups. Mothers that were seropositive for Chagas disease are depicted as M+, while B+ corresponds to babies diagnosed positive for cCD, and B-to babies diagnosed as non-infected. Thus, transmitting placentas come from M+B+ pairs, non-transmitting ones come from M+B-, and negative controls from M-B-. B0 refers to babies that did not complete the diagnostic algorithm.

The placental samples were selected and processed based on the characteristics of the clinical cases, meaning that the transmitting placentas corresponded to M+B+, the non-transmitting to M+B- and the controls to M-B-dyads. The experimental placental groups were set keeping the tightest equivalence in terms of type of delivery and newborn’s sex, including RIN values of the purified RNAs and the clinical characteristics described in Table 1.

**Table 1.** Clinical characteristics of the placental RNA samples used in the transcriptomic analysis.

| ID | Clinical group | Mother's age (years) | IgG Toxo | Gestation (weeks) | Pregnancy complications | Delivery | Newborn's sex | RIN RNA |
| --- | --- | --- | --- | --- | --- | --- | --- | --- |
| H014 | M-B- | 20 | Positive | 40 | PE | Cesarean | Male | 5.4 |
| H020 | M-B- | 26 | Negative | 39 | No | Vaginal | Male | 4.2 |
| M076 | M-B- | 25 | Negative | 40 | No | Vaginal | Male | 4.4 |
| M038 | M-B- | 39 | Negative | 39 | No | Vaginal | Female | 4.9 |
| 21NH | M-B- | 21 | Positive | 39 | No | Vaginal | Male | 6.1 |
| H013 | M-B- | 25 | Negative | 38 | No | Cesarean | Female | 5.4 |
| H011 | M-B- | 22 | Positive | 39 | No | Cesarean | Female | 5.1 |
| **M059** | M+B- | 30 | Negative | 40 | No | Vaginal | Male | 4.7 |
| **M108** | M+B- | 26 | Negative | 36 | PRM | Vaginal | Male | 4 |
| **M084** | M+B- | 35 | Positive | 37 | PE | Cesarean | Female | 3.2 |
| **48QP** | M+B- | 37 | Negative | 38 | No | Cesarean | Female | 4.6 |
| **M083** | M+B+ | 43 | Positive | 37 | PE | Cesarean | Male | 5.3 |
| **M030** | M+B+ | 25 | Positive | 38 | PE | Cesarean | Male | 4.7 |
| **M072** | M+B+ | 25 | Negative | 40 | Meco | Vaginal | Male | 4.5 |
| **15II** | M+B+ | 31 | Positive | 40 | No | Vaginal | Female | 4.9 |
| **M070** | M+B+ | 41 | Positive | 40 | No | Cesarean | Female | 4.3 |
| **M098** | M+B+ | 28 | Negative | 34 | PRM | Cesarean | Female | 4.1 |

ID: Patient identifier code. IgG Toxo: the mother's diagnosis of *Toxoplasma gondii* by indirect immunofluorescence assay. Results are reported as Positive or Negative according to a cut-off titer of 1:16. Titers $\geq 1:16$ were considered positive. PE: Preeclampsia. PRM: Premature Rupture of Membranes. Meco: Meconium.

The transcriptional profile of the placental samples was analyzed using a sample distance heatmap and its associated dendrogram. Since sample M098 was identified as an outlier based on its distance from all other samples, it was excluded from further analysis (S1 Fig).

### Differentially expressed genes (DEGs)

Differential gene expression analysis between all clinical groups (M+B+ vs. M-B-, M+B- vs. M-B-, and M+B+ vs. M+B-) is shown in Fig 2 and Tables 2-4. In each table, the gene identification (GID), the base 2 logarithm of the fold change (Log_2_FC), and the adjusted *p*-value (*p*_adj_) are shown. Only genes with a *p*_adj_ value ≤0.05 and |FC| > 1 are listed.

**Fig 2.**
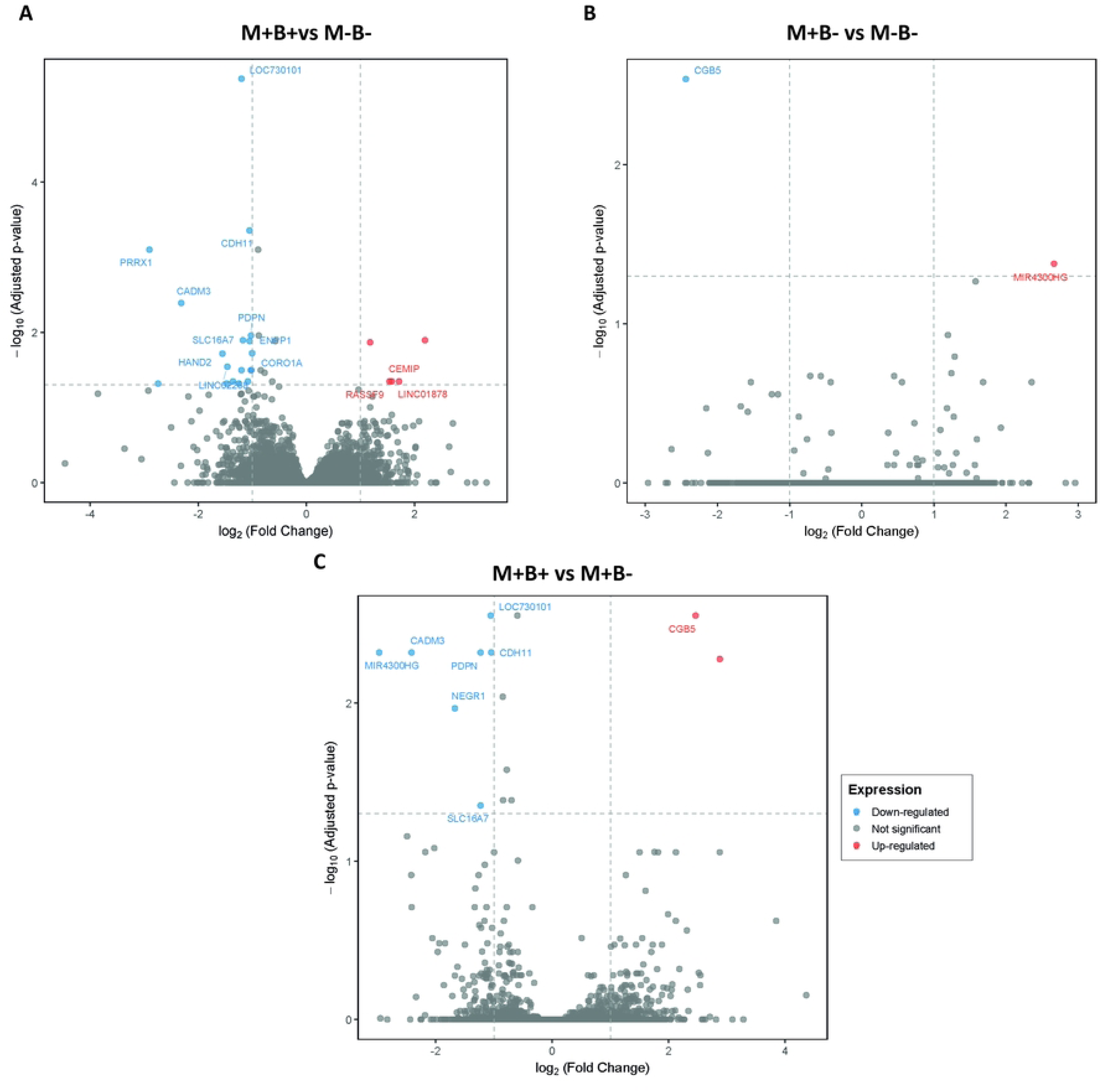
Differentially expressed genes. Volcano plots representing DEGs of M+B+ vs. M-B- (A), M+B-vs. M-B- (B), and M+B+ vs. M+B- (C). The Y-axis represents the negative logarithm of the *p*_adj_. The dashed line on the Y-axis indicates *p*_adj_ = 0.05. The X-axis shows log_2_FC and its dashed lines indicate 1 and -1. Red and blue dots represent the genes that were differentially expressed with a *p*_adj_ of 0.05 and | |FC| >1, and gray dots represent genes that did not surpass this cutoff.

**Table 2.**
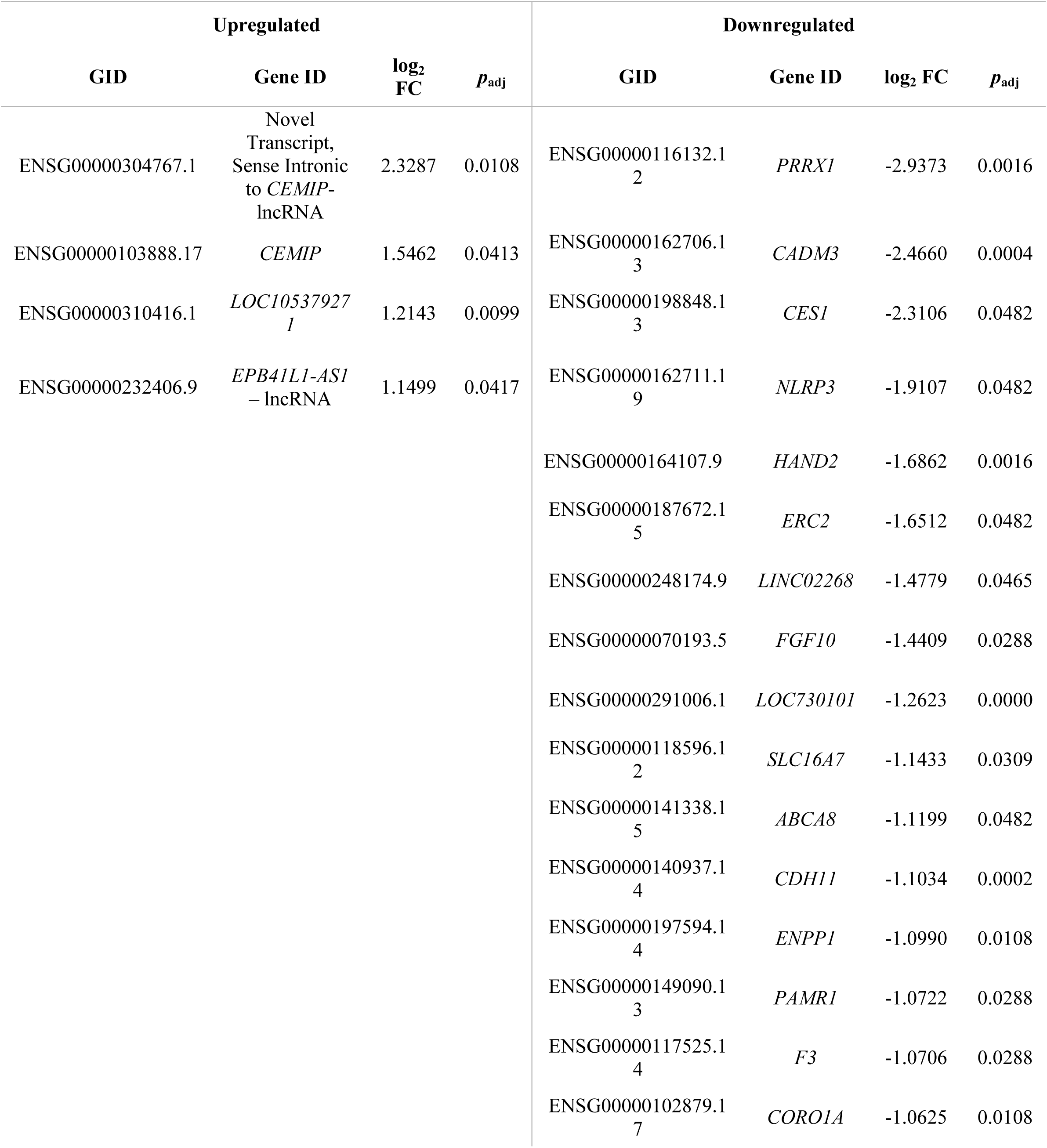
Differentially expressed genes in the M+B+ vs. M-B-comparison.

**Table 3.** Differentially expressed genes in the M+B-vs. M-B-comparison.

| Upregulated |  |  |  | Downregulated |  |  |  |
| --- | --- | --- | --- | --- | --- | --- | --- |
| GID | Gene ID | log<sub>2</sub> FC | p<sub>adj</sub> | GID | Gene ID | log<sub>2</sub> FC | p<sub>adj</sub> |
| ENSG00000245832.8 | MIR4300HG | 2.6632 | 0.0499 | ENSG00000189052.7 | CGB5 | -2.4383 | 0.0029 |

**Table 4.**
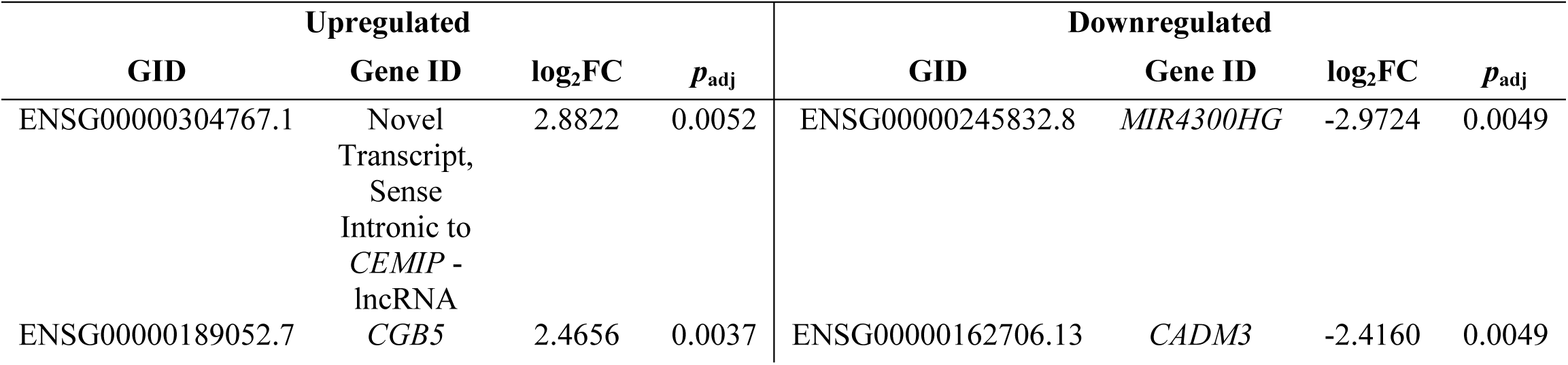
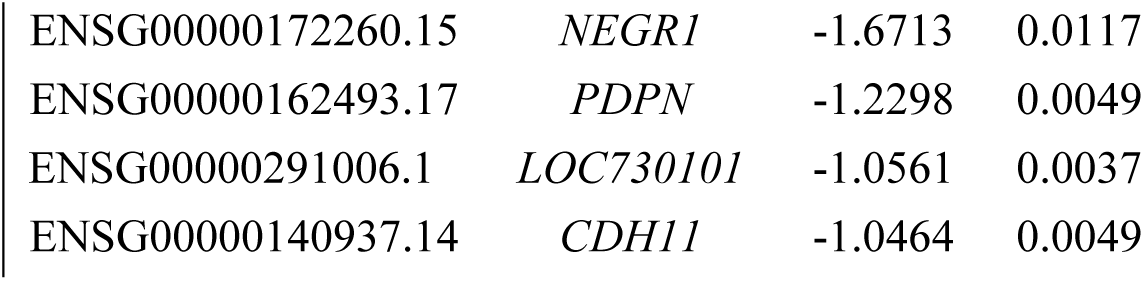
Differentially expressed genes in the M+B+ vs. M+B-comparison.

| Upregulated |  |  |  | Downregulated |  |  |  |
| --- | --- | --- | --- | --- | --- | --- | --- |
| GID | Gene ID | log<sub>2</sub>FC | p<sub>adj</sub> | GID | Gene ID | log<sub>2</sub>FC | p<sub>adj</sub> |
| ENSG00000304767.1 | Novel Transcript, Sense Intronic to *CEMIP* - lncRNA | 2.8822 | 0.0052 | ENSG00000245832.8 | MIR4300HG | -2.9724 | 0.0049 |
| ENSG00000189052.7 | *CGB5* | 2.4656 | 0.0037 | ENSG00000162706.13 | *CADM3* | -2.4160 | 0.0049 |

|  |  |  |  |
| --- | --- | --- | --- |
| ENSG00000172260.15 | *NEGR1* | -1.6713 | 0.0117 |
| ENSG00000162493.17 | *PDPN* | -1.2298 | 0.0049 |
| ENSG00000291006.1 | *LOC730101* | -1.0561 | 0.0037 |
| ENSG00000140937.14 | *CDH11* | -1.0464 | 0.0049 |

In the first contrast between transmitting (M+B+) and control placentas (M-B-) (Fig 2, Table 2), a total of 20 DEGs were identified, of which 16 were downregulated and 4 upregulated. Among the latter, two corresponded to long non-coding RNAs and one to an uncharacterized gene. The remaining upregulated gene was *CEMIP* (cell migration-inducing hyaluronidase 1), a protein involved in hyaluronan catabolism and in the positive regulation of protein phosphorylation and intracellular transport processes, likely mediated through its association with clathrin-coated vesicles during endocytosis.

Surprisingly, most of the DEGs were downregulated in transmitting placentas, and the highest negative fold-change corresponded to *PRRX1* (paired related homeobox 1), a transcriptional coactivator implicated in cell growth and differentiation. Other downregulated genes were those involved in cell adhesion, such as *CADM3* (cell adhesion molecule 3), which mediates both homophilic and heterophilic interactions, and *CDH11* (cadherin 11), a calcium-dependent transmembrane adhesion protein. Genes associated with inflammatory response regulation were also suppressed, including *NLRP3* (NLR family pyrin domain containing 3), a key component of the inflammasome complex that interacts with ASC/PYCARD, and *F3* (coagulation factor III), a membrane-bound glycoprotein that initiates coagulation cascades. In addition, genes involved in trophoblast invasion, proliferation and syncytialization were downregulated, such as *HAND2* (heart and neural crest derivatives expressed 2), a basic helix-loop-helix transcription factor, and *FGF10* (fibroblast growth factor 10), which plays critical roles in embryonic development, morphogenesis, and tissue repair.

In contrast, differential gene expression analysis between the M+B− and M−B− groups identified only two significantly altered transcripts (Fig 2, Table 3). *MIR4300*, a long non-coding RNA was upregulated and *CGB5* (chorionic gonadotropin subunit beta 5) was downregulated. *CGB5* encodes a protein produced by trophoblastic cells that plays a key role in stimulating ovarian steroidogenesis.

Subsequently, the comparison between transmitting (M+B+) vs. non-transmitting placentas (M+B-) (Fig 2, Table 4), showed 8 differentially expressed genes: 2 upregulated and 6 downregulated. The upregulated transcripts included a novel long non-coding RNA located within the intronic region of *CEMIP*, and *CGB5*, which encodes the chorionic gonadotropin subunit beta 5, previously noted for its role in trophoblast function.

Among the downregulated genes, several are associated with cell adhesion processes. These include *NEGR1* (neuronal growth regulator 1), predicted to localize to the extracellular region and plasma membrane; *PDPN* (podoplanin), a cell-surface glycoprotein involved in cell adhesion, chemotaxis, and the maturation of lymphatic and vascular structures; as well as *CDH11* and *CADM3*, previously described.

### Functional Enrichment Analysis

Gene set enrichment analysis (GSEA) using the gene ontology library (GO) between transmitting (M+B+) and control placentas (Fig 3A, S2 Table) revealed predominantly downregulated biological processes, particularly those involved in extracellular matrix organization, cell migration, and inflammation linked to chemotaxis. Similarly, in the non-transmitting (M+B-) vs. control comparison (Fig 3B, S3 Table), negatively enriched pathways were also dominant, including gene sets related to innate immune defense, inflammatory regulation, and pathogen recognition. Finally, contrasting transmitting vs. non-transmitting placentas (Fig 3C, S4 Table) showed positively enriched gene sets primarily associated with antigen presentation, immune regulation, and intracellular processes such as ER-Golgi vesicular transport and nucleosomal DNA organization. In contrast, underrepresented pathways were linked to embryonic development, extracellular matrix dynamics (e.g., collagen fibril assembly and basement membrane formation), and innate immune functions such as macrophage chemotaxis, phagosome maturation, and NK cell differentiation.

**Fig 3.**
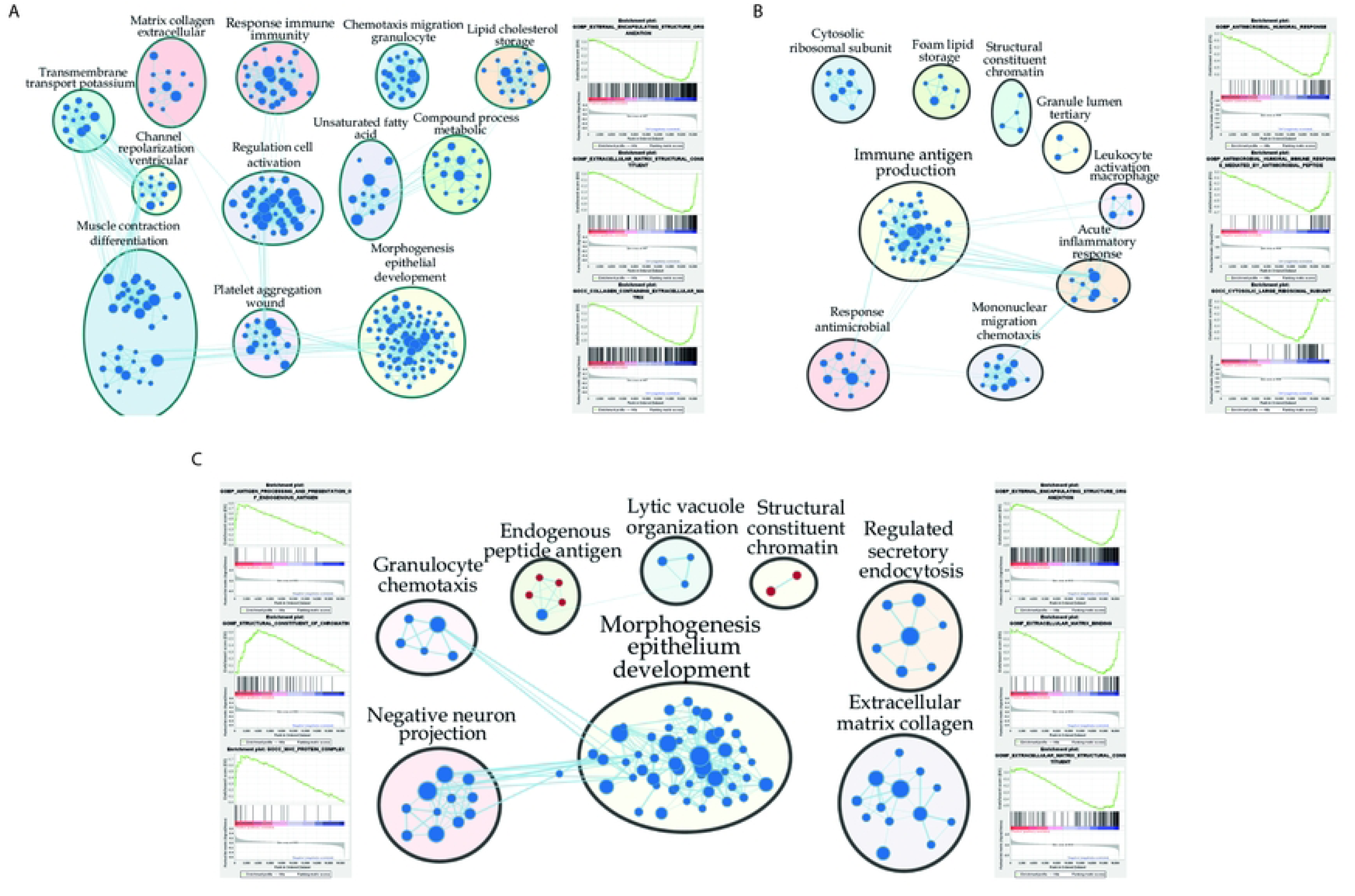
Gene set enrichment analysis: Gene Ontology library. Enrichment maps of M+B+ vs. M-B-(A), M+B-vs. M-B-(B), and M+B+ vs. M+B-(C). The diagrams show gene sets of altered GO terms with an FDR ≤ 10%. The color of the nodes represents the normalized enrichment score (NES), with red representing positive enrichment and blue representing negative enrichment. Related gene clusters are connected by light blue lines. GSEA results were visualized using Cytoscape. Enrichment plots for some of the most altered biological processes are shown on the sides.

GSEA using the Cell Type library was conducted to infer the cell types active in the different clinical groups (Fig 4, S5–7 Tables). Notably, gene sets associated with extravillous trophoblasts were overrepresented in the transmitting placentas compared to the other groups, but they were depleted in non-transmitting vs. control comparisons. An opposite pattern was observed for genes linked to villous trophoblasts and syncytiotrophoblasts: they were overrepresented in non-transmitting vs. control but depleted in transmitting vs. non-transmitting placentas.

**Fig 4.**
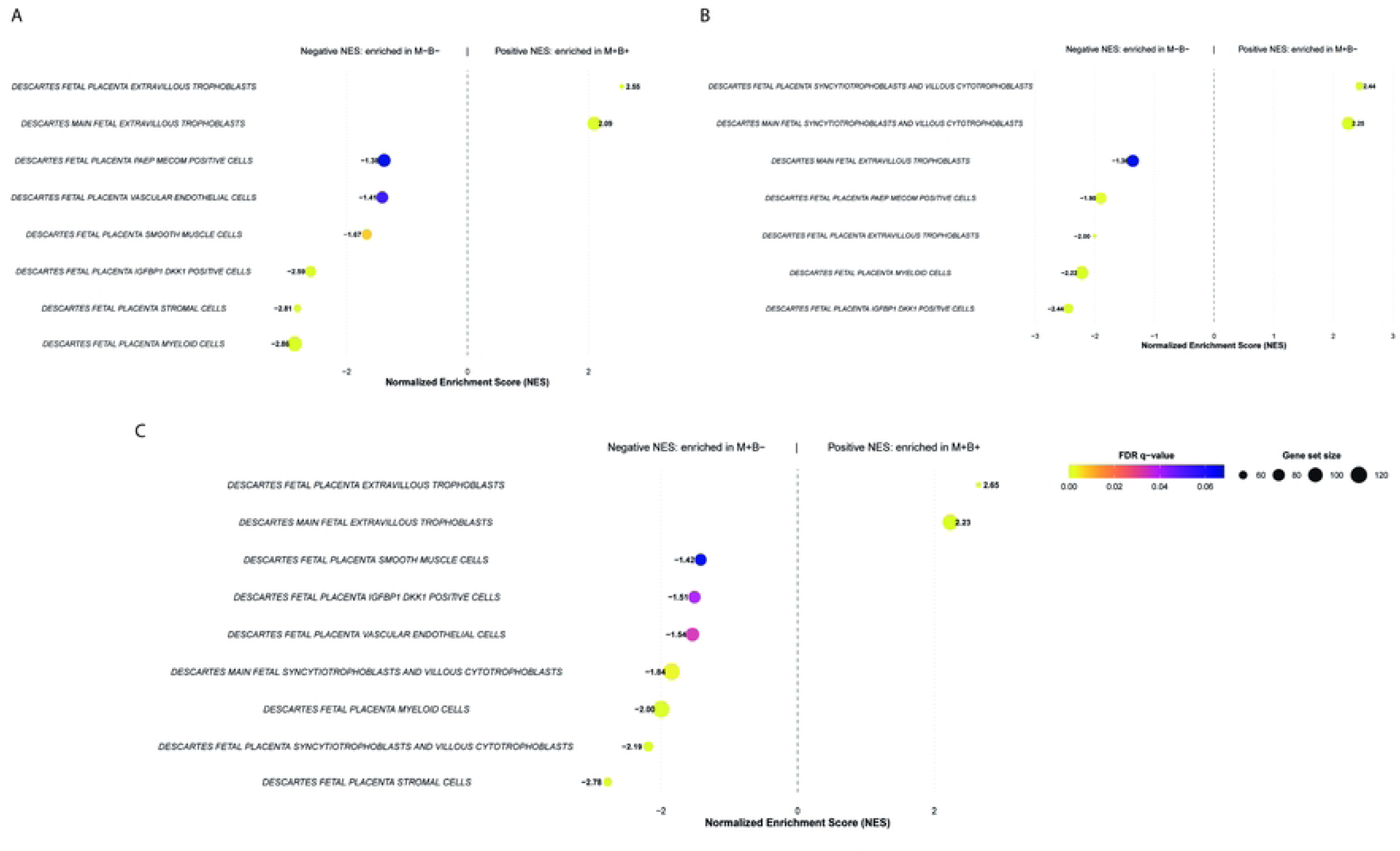
Gene set enrichment analysis: Cell type library. Dot plots of M+B+ vs. M-B-(A), M+B-vs. M-B-(B), and M+B+ vs. M+B-(C). The charts show gene sets of altered “cell type” terms with a False Discovery Rate (FDR) ≤ 10% related to placental groups. The normalized enrichment score is represented on the x-axis. FDR values closest to 0 are shown in yellow, blue those closest to 0.1, and size of the dot represents the size of the gene set (from 60 to 120).

Finally, when compared to the uninfected control, both *T. cruzi*-infected groups showed underrepresented cell types directly related to the placental immune system, regardless of infection transmission. Enrichment was observed in gene sets associated with fetal placenta myeloid cells and in PAEP⁺/MECOM⁺ and IGFB1⁺/DKK1⁺ cell populations.

### Gene co-expression network analysis

Co-expression network analysis identified three key modules—designated as dark grey, sky blue, and midnight blue— that displayed differential associations across clinical groups, namely a strong positive correlation with the control group and a strong negative correlation with the transmitting group (Fig 5A). This pattern suggests opposite transcriptional behavior of the same gene sets depending on the clinical phenotype. Among these, the dark grey module was selected for in-depth analysis due to two reasons. First, a large proportion of its genes were downregulated in the differential expression analysis comparing transmitting vs. control groups, consistent with the negative correlation observed in the co-expression network. Second, functional enrichment analysis of the dark grey module revealed a significant overrepresentation of genes associated with extracellular matrix organization, glycosaminoglycan binding, and growth factor interactions (Fig 5B). These findings suggest potential disruptions in cell– matrix communication and structural integrity in cases of cCD transmission.

**Fig 5.**
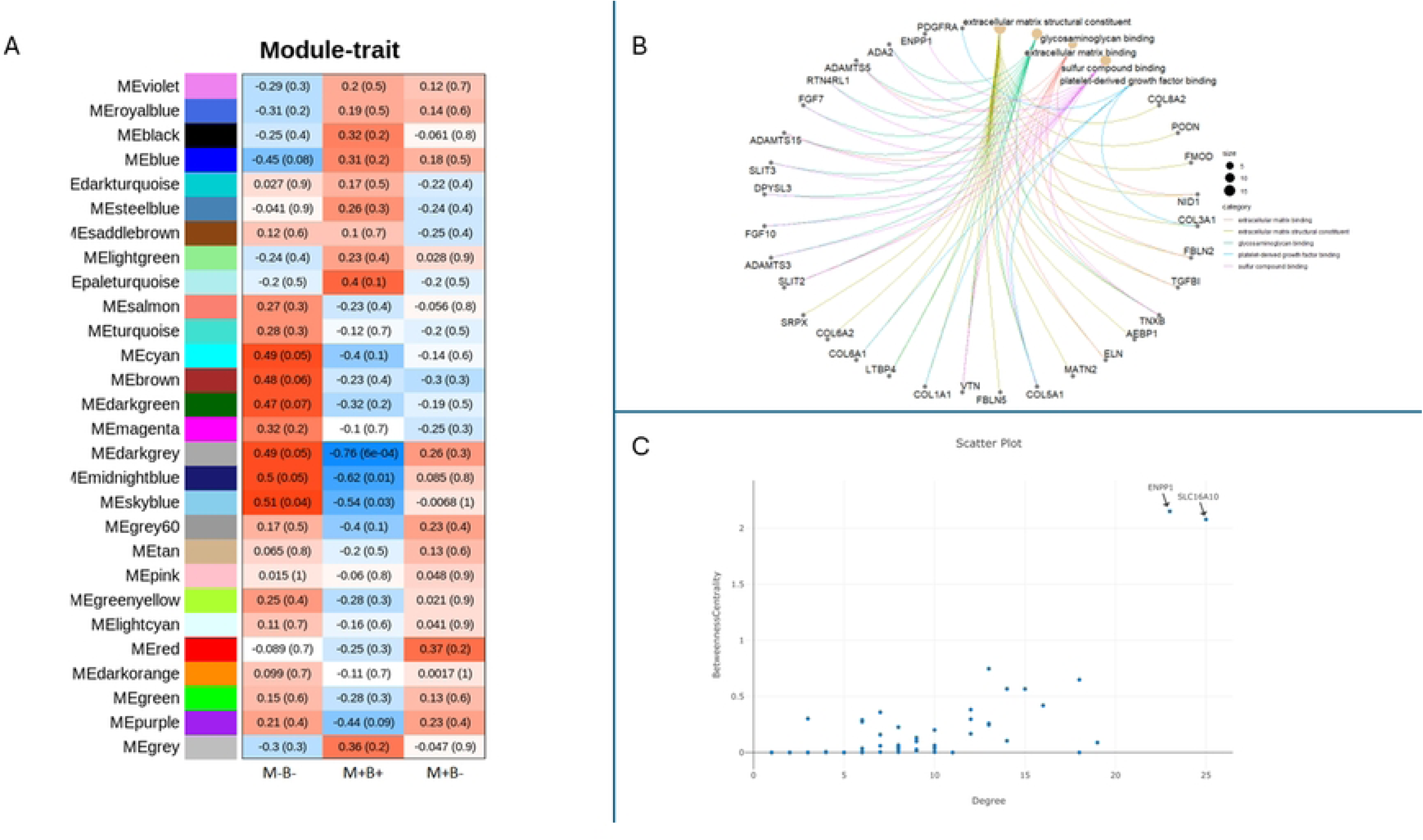
Network analysis of the dark grey module associated with the MB clinical groups. (A) Heatmap of module–trait relationships from WGCNA. (B) Functional enrichment of genes from the dark grey module. (C) Hub gene analysis within the module. The circular plot shows functional connections, and the scatter plot highlights key hub genes with high degree of betweenness centrality.

Moreover, two hub genes, *ENPP1* and *SLC16A10*, were identified based on centrality metrics (degree and betweenness centrality) within this module (Fig 5C). As central nodes in the co-expression network, these genes may play pivotal roles in the biological pathways altered during congenital infection. Briefly, *ENPP1* (ectonucleotide pyrophosphatase/phosphodiesterase 1) is a transmembrane glycoprotein, member of the ecto-nucleotide pyrophosphatase/phosphodiesterase (ENPP) family, while *SLC16A10* -solute carrier family 16 member 10, belongs to a family of membrane transporters that mediate Na(+) transport of aromatic amino acids across the cell membrane.

## Discussion

Although the incidence of vertical transmission of *T. cruzi* is relatively low [11], it remains the main transmission route due to successful vectorial control, transfusion and transplant-related cases [12,13]. Congenital CD has declined over recent decades, particularly in the areas of the present study. In Chaco Province, the incidence in the period 2014–2015 was 7.1% at Hospital Perrando [14], while in Salta Province in 2021 it was approximately 1.6% (according to non-published data from Hospital Público Materno Infantil). However, completing the diagnostic algorithm presents challenges for the vulnerable affected population, as demonstrated in the present study where in 35.8% newborns this algorithm was not completed, leading to their exclusion—an issue previously reported and further exacerbated by the COVID-19 pandemic [15–17]. Consequently, it was necessary to enroll 162 *T. cruzi*–infected pregnant women to obtain 12 placental samples from confirmed congenital transmission cases.

The human placenta is a highly specialized organ whose structure and function rely on tightly regulated processes, including continuous syncytiotrophoblast turnover, vascular remodeling by extravillous trophoblasts, and finely tuned immune regulation to prevent fetal rejection and protect it against infection. These coordinated processes result in a complex, cell type–specific transcriptional landscape [18].

### Differentially expressed genes

#### M+B+ vs. M-B-

This comparison revealed 20 differentially expressed transcripts, including four upregulated and 16 downregulated genes. Notably, a quarter of these transcripts were non-coding RNAs (ncRNAs), emphasizing their emerging relevance in placental gene regulation. This observation aligns with recent findings that implicate lncRNAs in key trophoblast functions, including proliferation, migration, invasion, and apoptosis [19,20]. Among the upregulated transcripts, two were particularly noteworthy: *CEMIP* (cell migration-inducing hyaluronidase 1) and a novel lncRNA (*ENSG00000304767*) located within an intron of the *CEMIP* gene. *CEMIP* is known to participate in extracellular matrix degradation and intracellular transport via clathrin-mediated endocytosis and has also been associated with trophoblast migration and invasiveness [21,22]. The co-upregulation of *CEMIP* and its intronic lncRNA suggests a potential regulatory relationship. Previous studies have shown that *CEMIP* expression can be modulated by lncRNAs such as *LINC00958*, *CASC19*, and *HCP5*, which act by sequestering specific miRNAs [23–25]. In this study, the intronic transcript *ENSG00000304767* was found within a genomic region marked by enhancer-associated epigenetic signatures and overlapping the predicted enhancer *ENSR15_BL2T2*, which may regulate *CEMIP* (S1 Fig2B). Its expression pattern and localization suggest that it could function as an enhancer RNA, modulating the transcriptional activity of *CEMIP* or nearby genes— though experimental validation is required to confirm this hypothesis.

The involvement of structural pathways in congenital transmission is supported by the fact that several genes associated with cell adhesion were downregulated. These included *CADM3* and *CDH11*. *CADM3* encodes a calcium-independent adhesion molecule capable of both homophilic and heterophilic interactions via nectin dimerization [26,27]. Although it has been linked to preeclampsia [28,29], its specific function in placental tissue remains underexplored. *CDH11*, a type II classical cadherin, plays a calcium-dependent role in cell–cell adhesion and is expressed in syncytiotrophoblasts and extravillous cytotrophoblasts. It contributes to the epithelial–mesenchymal transition required for anchoring to the decidua [30,31]. Notably, *CDH11* expression increases during trophoblast syncytialization, and its overexpression has been shown to suppress proliferation and enhance differentiation [32].

Among the genes analyzed in this contrast, *PRRX1* (paired related homeobox 1) exhibited the highest downregulation. The *PRRX1* protein acts as a transcriptional co-activator, boosting the DNA-binding capacity of serum response factor, which in turn activates genes in response to growth and differentiation signals. *PRRX1* has also been shown to promote the expression of *TGFβ3* and fibrillar collagen genes [33]. Recently, *PRRX1* was identified as a transcriptional regulator of mRNA expression of *COL6A3* (collagen type VI alpha 3 chain) in adipose cells [34]. Therefore, *PRRX1* function is closely related to the extracellular matrix.

Evidence from both *in vitro* studies [35–37]) and histopathological analyses of placentas from seropositive women [38] indicates that *T. cruzi* infection induces placental tissue damage, including syncytiotrophoblast detachment, which may facilitate parasite invasion. Other studies reported destruction of the syncytiotrophoblast and villous stroma, along with selective disruption of the basal lamina and collagen I organization [39]. The gene expression changes identified in our study indicate the occurrence of trophoblast detachment and extracellular matrix remodeling in placental tissues associated with congenital transmission. Specifically, downregulation of genes involved in trophoblast function, such as *CEMIP*, *HAND2* and *FGF10*, suggest impairment of epithelial turnover and extravillous trophoblast proliferation and invasion —the main defense mechanisms reported to protect against infection [2,40]. *HAND2* encodes a transcription factor that has been implicated in trophoblast invasion, proliferation, and syncytialization [41–43], in addition to its role in regulating the timing and progression of parturition [44,45]. In fact, dysregulation of *HAND2* expression or signaling has been associated with preterm birth and preeclampsia [46,47].

The long intergenic non-protein coding RNA 2268, *LINC02268 (ENSG00000248174)* has been reported to co-express itself with other genes and is potentially involved in the activation of IL-6/JAK/STAT3 pathways in cancer [48]. GeneCards predictions suggest it could regulate *HAND2* (among others), which is particularly interesting since *HAND2* also appeared downregulated in our study (Table 2).

Finally, downregulated *NLRP3* and *F3* evidenced an altered immune status compared to controls. Notably, the inflammasome activation by *NLRP3* has been reported as essential to control *Leishmania*, *T. gondii* and *T. cruzi* infections. On the contrary, inactivation of this pathway inhibits the differentiation of T-cell responses and killing of *T. cruzi* by macrophages, resulting in greater parasite burdens [49]. Additionally, parasite– macrophage interaction may trigger TF release, promoting coagulation, inflammation, and frequent thrombotic events in CD patients [50]. In our study, the low expression of *NLRP3* suggests limited inflammation, which in turn would favor the development of the infection and consequently congenital transmission.

#### M+B-vs. M-B-

Only two DEGs between non-transmitting and control groups were detected: *MIR4300HG*, a long non-coding RNA that hosts *MIR4300*, was upregulated (FC = 2.66), and *CGB5* was downregulated. Although the function of *MIR4300* remains unclear, it is known that microRNAs broadly regulate gene expression and biological processes [51], and while the mature microRNA was not directly detectable due to its small size, its expression is potentially occurring. Using miRWalk 3.0, *CGB5* was predicted as a potential *MIR4300* target, with binding sites in its coding region, suggesting possible post-transcriptional repression. In turn, *CGB5* encodes the β-subunit of human chorionic gonadotropin (β-hCG), a hormone predominantly expressed in trophoblasts through *CGB3*, *CGB5*, or *CGB8* [52]. Its underexpression is consistent with findings from Juiz et al. [10](2018) in non-transmitting placentas. It has been shown that β-hCG production was increased in human placental chorionic villus explants and *in vitro* using the human trophoblast cell line BeWo infected with the Ypsilon strain (TC II) [40,53]. In contrast, human chorionic villi explants challenged with an isolate from a cCD case (Lucky, TcII/VI) and the Tulahuen strain (Tc VI) showed a decrease in β-hCG production [54]. Since β-hCG is considered an indicator of the vitality of syncytiotrophoblast [55], our findings suggest a functional alteration of the placental barrier.

#### M+B+ vs. M+B-

This comparison revealed that in the transmitting group the long non-coding RNA intronic to *CEMIP* and *CGB5* were overexpressed, while the genes downregulated were *MIR4300HG*, *CADM3*, *NEGR1*, *PDPN*, *LOC730101*, and *CDH11*. As previously mentioned, *CADM3* is related to cell adhesion, as well as *NEGR1* and *PDPN*, while *CDH11* is involved in trophoblast differentiation. *NEGR1* is a membrane-bound protein involved in intercellular adhesion, widely studied in the brain, but also expressed in the placenta, where its specific function remains unknown [56]. Podoplanin, *PDPN*, is a cell-surface receptor implicated in cell adhesion, chemotaxis, and lymphatic and vascular maturation. In human placentas, it is mainly expressed in decidual cells and chorionic villous stromal cells, particularly during pregnancy [57]. Interestingly, the authors propose that low levels of podoplanin could cause inflammation in the villi, at least in the context of molar pregnancies [57]. These results suggest that the main differences between parasite transmission and non-transmission in infected placentas rely on trophoblast differentiation and functionality, cell adhesion, and the extracellular matrix integrity.

### Gene set enrichment analysis

#### M+B+ vs. M-B-

In this comparison, most downregulated terms in the transmitting group are related to the extracellular matrix, particularly collagen organization and metabolism. As previously mentioned, when describing the downregulated DEGs in the same group, cell adhesion and extracellular matrix organization processes are closely linked to trophoblast detachment and extracellular matrix reorganization in response to *T. cruzi* infection [35–39].

Gene sets associated with metallopeptidase activity—enzymes that degrade proteins through a metal ion in their active site—were also found to be downregulated. This finding is particularly noteworthy, as it supports previous evidence linking two MMP2 gene polymorphisms with the occurrence of cCD [58]. Polymorphisms that reduce metalloproteinase transcription may impair the placental response to infection, given the role of these enzymes in extracellular matrix degradation, epitope processing and modulation of immune and inflammatory responses [59,60]. Our study suggests reduced leukocyte migration by the enrichment of terms such as GOBP_LEUKOCYTE_CHEMOTAXIS and GOBP_MYELOID_LEUKOCYTE_MIGRATION. In turn, this impairment may be caused by decreased metalloproteinase activity and extracellular matrix remodeling, as effective leukocyte infiltration requires both matrix degradation and the generation of chemotactic signals derived from metalloproteinase-mediated proteolysis [59]. Finally, placentas from transmitting mothers showed an overall downregulation of the immune response compared to controls. Several gene sets related to innate and adaptive immune responses had negative NES values, but those related to the inflammatory response were particularly diminished, potentially facilitating placental transmission of the parasite [61].

#### M+B-vs. M-B-

GSEA comparing these groups revealed a downregulation of both innate and adaptive immune responses. The most prominently affected gene sets were related to the humoral immune response, particularly those mediated by antimicrobial peptides, which overlapped with the signatures previously reported by Juiz et al. [10](2018). However, in contrast to Juiz et al., who reported a general upregulation of immune responses in non-transmitting cases, our findings indicate a downregulation of these responses in the same group. Transcriptomic changes were evaluated in *in vitro* assays using human chorionic villus explants infected with *T. cruzi* Y strain (DTU II) at different parasite concentrations and incubation times [62]. Modulation of gene expression was dependent on both factors, with most DEGs downregulated in three of the four tested conditions. The most pronounced transcriptional changes occurred after 24 hours of exposure to the lowest parasite concentration. In both the present study and the *in vitro* model, the immune response emerged as the most affected pathway; however, while it was consistently downregulated in our data, it was variably up-or downregulated depending on infection conditions in the explant model [62].

#### M+B+ vs. M+B-

Finally, direct comparison of placental transcriptomes from M+B+ and M+B− revealed increased antigen processing and presentation in transmitting placentas, while extracellular matrix organization and collagen-related processes were reduced. Cell type enrichment analysis indicated decreased syncytiotrophoblast and cytotrophoblast signatures in transmitting cases, suggesting impaired epithelial renewal, a process thought to protect against vertical transmission [2]. Additionally, gene sets related to development and morphogenesis were significantly downregulated in the transmitting group. This aligns with clinical observations of shorter gestation in infected newborns as well as low birth weight, Apgar scores <7, and increased rates of premature membrane rupture in babies born to seropositive mothers compared to controls [63]. GSEA using the Cell Type library identified differential expression patterns in villous, extravillous and syncytiotrophoblasts, suggesting that shifts in specific trophoblast populations may be associated with susceptibility or resistance to transplacental transmission of *T. cruzi* (Fig 4, S5-7 Tables).

## Gene co-expression network analysis

Three key modules (e.g., dark grey, sky blue and midnight blue) showed correlations with clinical groups. Among them, dark grey was prioritized for further analysis because it showed the strongest concordance between co-expression patterns, differential expression and biological function. Notably, the eigengene expression of this module exhibited an inverse correlation with the occurrence of *T. cruzi* vertical transmission, indicating that genes within this module are upregulated in uninfected controls and downregulated in cases of cCD [64,80]. Furthermore, a substantial proportion of DEGs between transmitting and control placentas (10 out of 16) overlapped with genes in this module. Collectively, these results suggest the existence of a coordinated transcriptional repression program linked to congenital *T. cruzi* transmission [8]. Additionally, functional enrichment analysis of the dark gray module revealed a significant overrepresentation of genes involved in extracellular matrix organization, glycosaminoglycan binding, and growth factor interactions, suggesting a disruption in cell-matrix communication and structural integrity in transmitting placentas, confirming the GSEA results [65]. Finally, two genes were identified as hub genes: *ENPP1* and *SLC16A10*. The protein encoded by *ENPP1* has recently been associated with the innate immune response, among other functions, and it has been suggested that its variants are linked to different disease phenotypes [66]. Similarly, downregulation of *SLC16A10* has been proposed to be linked to gestational disorders [67]. These 2 genes deserve further investigation as potential biomarkers of transmission risk, especially taking into account that *ENPP1* was also significantly downregulated in the DEGs analysis.

## Conclusions

This transcriptomic study is the first to compare placental tissues from *T. cruzi*-infected mothers who transmitted congenital Chagas disease (CCD), those who did not, and non-infected controls. Through an integrated analytical strategy, we identified both coding and non-coding genes with potential regulatory functions—most notably *CEMIP* and the intronic long non-coding RNA *ENSG00000304767*—alongside several genes implicated in cell adhesion, suggesting a possible compromise of the placental barrier integrity during transmission. GSEA revealed distinct functional alterations across groups, with transmission cases showing marked changes in extracellular matrix-related pathways and in key trophoblast cell types. These findings were further confirmed by co-expression network analysis, which highlighted *ENPP1* and *SLC16A10* as hub genes potentially involved in vertical transmission mechanisms. Together, these results provide new molecular insights into the biology of placental tissue in congenital *T. cruzi* transmission and offer promising candidates for future functional and translational studies.

## Materials and methods

### Ethical statement

The study was conducted upon approval of the bioethics committees of the participating institutions, following the principles of the Helsinki declaration, in accordance with resolution 1480/2011 of the Ministerio de Salud from Argentina. Patients were recruited at the Hospital Perrando, Resistencia, Province of Chaco and at the Hospital Público Materno Infantil, Salta, Province of Salta. In all cases, the purpose of the study was explained to the mothers, and informed consent was obtained before sample collection.

### Subjects and samples

Pregnant women were diagnosed with CD using conventional serological methods performed at their respective healthcare centers as part of routine screening. Standard clinical guidelines also included screening for toxoplasmosis [68], preeclampsia [69], premature rupture of membranes [70] and meconium-stained amniotic fluid.

Only infants born to seropositive mothers were tested for CCD, following the current diagnostic algorithm. This includes a micromethod assay and/or qPCR to detect parasitemia within the first days of life or between 4 and 8 weeks of age, followed by confirmatory serological testing—using two distinct serological methods—after the infant reaches nine months of age [71].

Fresh normal placentas were obtained immediately after labor from vaginal or cesarean deliveries, kept at 4 °C and processed within the next 3 h. Subsequently, each placenta was dissected: a portion of tissue located at 4 cm from the umbilical cord was extracted, and the middle region was immersed in RNAlater Solution (Applied Biosystems, Foster City, CA), overnight at 4 °C to facilitate tissue penetration. Finally, the samples were stored at -80 °C until RNA extraction.

### RNA extraction

Placental samples preserved in RNA-Later Solution were thawed and always kept cold. Between 50 mg and 100 mg of placental tissue were cut and weighed on a Petri dish. The RNeasy Lipid Tissue Mini Kit (Qiagen, Hilden, Alemania) was used according to the manufacturer’s instructions. The tissue was disaggregated after adding 1 mL of Qiazol, using the Fisherbrand™ Pellet Pestle™ Cordless Motor with clean Pellet Pestles (Thermo Fisher Scientific, Waltham, Massachusetts, USA). DNA digestion was also performed using an RNase-Free DNase Set (Qiagen) on the same column according to the manufacturer’s instructions. Elution was performed with 50 μl of DEPC-treated water. Aliquots were separated from the RNA obtained to have a first approximation of the concentration; purity of the RNA was assessed using the DS-11 Spectrophotometer (DeNovix, Wilmington, USA) and integrity was verified after electrophoresis in 1% Agarose Gel, with 0.5X TBE buffer. The remaining RNA solution was precipitated by adding 0.1 volumes of Sodium Acetate, pH 5.5, mixing gently and then adding 2 volumes of 100% ethanol and mixing again. The precipitated RNA was stored at -80°C until shipping.

### Transcriptomic studies

An RNA-seq study was performed using samples from three clinical groups independently analyzed: control or uninfected placentas (M-B-, n=7), infected but non-transmitting placentas (negative for cCD or M+B-, n=4), and infected, transmitting placentas (positive for cCD or M+B+, n=5). The preparation of cDNA libraries and sequencing was performed by Macrogen (Seoul, South Korea): samples were used individually to construct the libraries using the TruSeq Stranded Total RNA with Ribo-Zero H/M/R_Gold kit (Illumina, San Diego, CA). Sequencing was performed using Illumina NovaSeq 6000 platform with a 101 bp paired-end reads strategy (Supplementary Table 1). The RNA-Seq data have been submitted to the National Center for Biotechnology Information (NCBI) Sequence Read Archive (SRA) under the BioProject accession number PRJNA1297177.

### RNA sequencing analysis

Raw read quality was initially assessed using FastQC (version 0.11.5; Babraham Bioinformatics, Cambridge, United Kingdom). Subsequently, adapter sequences were removed using Trimmomatic, version 0.36, in paired-end mode [72], and the quality of the trimmed reads was re-evaluated with FastQC. The cleaned reads were then aligned to the human reference genome GRCh38.p14, obtained from Gencode (https://www.gencodegenes.org/human/), using STAR, version 2.5.2b [73] with default parameters, generating coordinate-sorted BAM files. The integrity of these BAM files was verified using Samtools, version 1.3.1, via its flagstat function [74].

For quantification, FeatureCounts, version 2.0.1, was employed in paired-end mode [75] to generate a count matrix, which was subsequently transformed into a DESeq2 object (dds). A filtering criterion [rowSums(counts(dds) ≥ 3) ≥ 6] was applied to retain only those genes with at least 3 counts in a minimum of 6 samples, thereby minimizing noise from lowly expressed genes. Finally, differential expression analysis was performed using the DESeq2 package, version 1.44.0, [76], facilitating the identification of differentially expressed genes. DEGs were identified using an adjusted p-value threshold of 0.05 and an absolute log₂ fold-change (|log₂FC|) greater than 1.

Functional enrichment analysis was conducted using the ClusterProfiler package, version 4.14.6, [77], applying thresholds of p-value Cutoff > 0.01 and q-value Cutoff > 0.05. P-value correction was performed using the Benjamini-Hochberg method, and the results were visualized with ggplot2. Considering that the hierarchical structure of Gene Ontology (GO) terms can lead to redundancy in the list of enriched terms, the simplify function from the GOSemSim package was employed with default thresholds to evaluate semantic similarities among GO terms, effectively eliminating closely related terms, while retaining only the most representative ones. Gene set enrichment analysis was performed using the GSEA software, version 4.3.3, [78] with the MSigDB gene sets [79], specifically the C5 collection (ontology gene sets, 16,107 gene sets) and the C8 collection (cell type signature gene sets, 840 gene sets), employing 1000 permutations and default settings for all other parameters. For GSEA analysis using the whole expression matrix, an FDR < 25% was considered significant.

### Co-expression gene analysis

Gene coexpression networks were constructed using the WGCNA package version 1.72-1, [80] on variance-stabilized DESeq2 version 1.44.0, [76] data. Genes with low expression were removed following the initial filtering criteria described above, and the expression matrix was further filtered to include only genes with variance in the upper 75th percentile to improve network robustness.

Similarities in gene expression between pairs were calculated using Pearson’s correlation coefficient. These were transformed into adjacency matrices using a soft-threshold of power 14, selected to approximate a scale-free topology according to standard WGCNA criteria [80]. A signed network was constructed, including only positive correlations (genes with similar expression patterns), while negative correlations were downweighted to better reflect biologically relevant coexpression. The topological overlap matrix (TOM) and its dissimilarity (dissTOM) were calculated to assess the interconnectedness of the network.

Modules of co-expressed genes were identified by hierarchical clustering of the dissTOM, followed by dynamic tree cutting (minimum module size = 30 genes). Modules with high eigengene similarity were merged based on a predefined soft-threshold.

Module eigengenes were correlated with clinical traits (M-B-, M+B-, M+B+) to identify phenotype-associated modules. Hub genes were defined as those with high node degree (the number of connections a gene has within the module), and high betweenness centrality (the extent to which a gene acts as a bridge between other genes) within the co-expression network. Functional annotation of selected modules was performed using ClusterProfiler package, version 4.14.6, [77], and networks were visualized with Cytoscape version 3.10.0, [81].

## Supporting information

## Acknowledgments

We extend our thanks to Patricio Yankilevich for his contribution to the preliminary bioinformatic analysis of the RNA-seq data. We are also deeply grateful to the healthcare teams at Hospital Perrando (Province of Chaco) and Hospital Público Materno Infantil (Province of Salta) for their dedication to the recruitment and follow-up of mothers and their newborns, whose placental samples made this research possible. Most importantly, we thank the mothers who generously agreed to participate in this study, with the shared hope that these findings may ultimately contribute to improving the health and well-being of their children.

## Funds

AGS received funds from MinCyT PICT 2020-0862 and PICT 2021-098.

PZ received funds from CONICET PIP 2021-3121 and MinCyT PICT and PUE 22920170100106CO.

